# Deep Microbial Colonization in 2-Billion-Year-Old Ultramafic Rock from the Bushveld Complex

**DOI:** 10.64898/2026.04.13.717956

**Authors:** Taro Kido, Susan J. Webb, Mariko Kouduka, Hiroki Suga, Hanae Kobayashi, Toshiaki Ina, Takahiro Kawai, Takanori Wakita, Takuma Kaneko, Tomoya Uruga, Masaki Oura, Julio Castillo, Jens Kallmeyer, Kgabo Moganedi, Amy J. Allwright, Reiner Klemd, Frederick Roelofse, Mabatho Mapiloko, Stuart J. Hill, Clement Ndou, Thulani Maupa, Lewis D. Ashwal, Robert B. Trumbull, Yohey Suzuki

## Abstract

Archean cratons may provide stable microbial habitats in the deep subsurface, as evidenced by the discovery of billion-year-old crustal fluids^1,2^. However, the long-term habitability of these cratonic environments is uncertain, as polymetamorphic evolution in most cratons typically destroys microbial habitats through mineral reactions and porosity loss^3,4^. Preservation of deep microbial habitats is more likely where mantle-derived magma intruded the craton after metamorphic overprinting^4^. Here we report the discovery of dense microbial colonization at 814 m depth within the 2.05-billion-year-old, unmetamorphosed Bushveld Igneous Complex intrusion, South Africa^5^. Using advanced contamination-control protocols^6,7^ and synchrotron-based X-ray spectroscopy, we identified indigenous microbial cells localized at the rims of phlogopite, a hydrous phyllosilicate mineral. Our study reveals that microbial colonization is associated with Fe(III) derived from the structure of phlogopite, where the dehydrogenation likely oxidizes Fe(II) to Fe(III) coupled to H_2_ generation^8^. Despite the absence of fracture-driven fluid ingress in the unfractured rock matrix, aqueous alteration evidenced at the rims by potassium removal indicates a self-sustaining habitat driven by an internal redox gradient^9^. These findings demonstrate that aqueous alteration of ultramafic rocks can sustain isolated microbial life over geological timescales, significantly expanding the potential for long-term habitability on both Earth and Mars^4,10^.

## Main

Archean cratons are old, stable components of the continental lithosphere that are preserved in many places around the globe^11^. Recently, it has been found that crustal fluids have remained isolated in deep cratonic rocks for 1.2 and 1.7 billion years, respectively, of the Kaapvaal Craton in South Africa^1^ and the Laurentia Craton in Canada^2^. Given that microbes and abiotically generated organic compounds (e.g., acetate and formic acid) have been found in isolated crustal fluids^12,13^, Archean cratons may therefore have provided stable microbial habitats throughout the history of life on Earth. Despite the abundance of energy sources such as H_2_, CH_4_, and organic acids^12,13^, microbial cell densities in these crustal fluids are extremely low^14,15^. This low cell density may result from the scarcity of dissolved electron acceptors, such as nitrate, sulfate, and dissolved inorganic carbon in the fluids^12,13^. Another possible explanation for this low cell density is that the rocks reveal evidence for multiple tectono-thermal metamorphic events^16^, which may have substantially reduced rock porosity and the bioavailability of solid-state oxidants^3^.

Following metamorphic overprinting, Archean cratons commonly experienced magmatic intrusions of mafic-ultramafic magmas from the underlying mantle^4^, which resulted in the formation of thick cumulate layers rich in olivine and pyroxene. The Rustenburg Layered Suite of the Bushveld Igneous Complex in the Kaapvaal Craton in South Africa is the largest known intrusion of this kind^5^, and it has not suffered significant metamorphic alteration since its emplacement in the Paleoproterozoic Transvaal Supergroup approximately 2.05 billion years ago (ca. 2.05 Ga)^17,18^. We studied a freshly-drilled rock core sample obtained from a depth of ca. 814 m by the Bushveld Complex Drilling Project (BVDP) of the International Continental Scientific Drilling Program (ICDP)^19^ at the Marula Platinum Mine, South Africa (24.50906°S, 30.08757°E).

### Microbial colonization in unfractured pyroxenite

In an earlier study of the BVDP, we collected a drill-core sample from a depth of ca. 15 m that was composed of norite (a rock dominated by plagioclase and orthopyroxene) and with abundant fractures to test on-site procedures to monitor drilling-fluid contamination using fluorescent microspheres with a diameter range from 0.25–0.45 μm, similar to the size of microbial cells^6,20^. We also established subsequent laboratory procedures for decontamination, for counting fluorescent microspheres and microbial cells, and for visualization within a rock section^6^. This sample contained microbial cells located around mineral-filled fractures that were free of fluorescent microspheres, showing they were uncontaminated.

We applied the same on-site and laboratory procedures to a study of the deeper sample (ca. 814-m), which was not intersected by fractures (Fig. 1a). Under ultraviolet (UV) illumination, the blue fluorescence of microspheres on the core surface was intense (Fig. 1b; Extended Data Fig. 1a), demonstrating proper delivery of the tracer to the bottom of the borehole through the circulating drilling fluid. The core was broken open with a sterilized hammer, and the fresh surface was observed under UV illumination and found to be partly contaminated at the edges only (Fig. 1c). The interior and exterior of the core were then separated using a sterilized rock trimmer. Fluorescent microspheres were abundant in the core exterior (4.9 ± 1.3 × 10^6^ microspheres cm^−3^) but significantly depleted in the core interior (3.1 ± 0.3 × 10^4^ microspheres cm^−3^). Given the concentrations of the microspheres (3.7 ± 1.8 × 10^8^ microspheres mL^−1^) and microbial cells (3.2 ± 0.1 × 10^6^ cells mL^−1^) in the drilling fluid, the number of contaminant microbial cells in the core interior is calculated to be less than 3.0 × 10^2^ cells cm^−3^. This low level of contamination demonstrated that the core interior was suitable for subsequent microbiological analysis. Petrographic observation in a thin section revealed that the sample was pyroxenite, with orthopyroxene and clinopyroxene contents of ca. 60% and ca. 30%, respectively, and with <10% combined phlogopite and quartz (Fig. 1d).

**Fig. 1.**
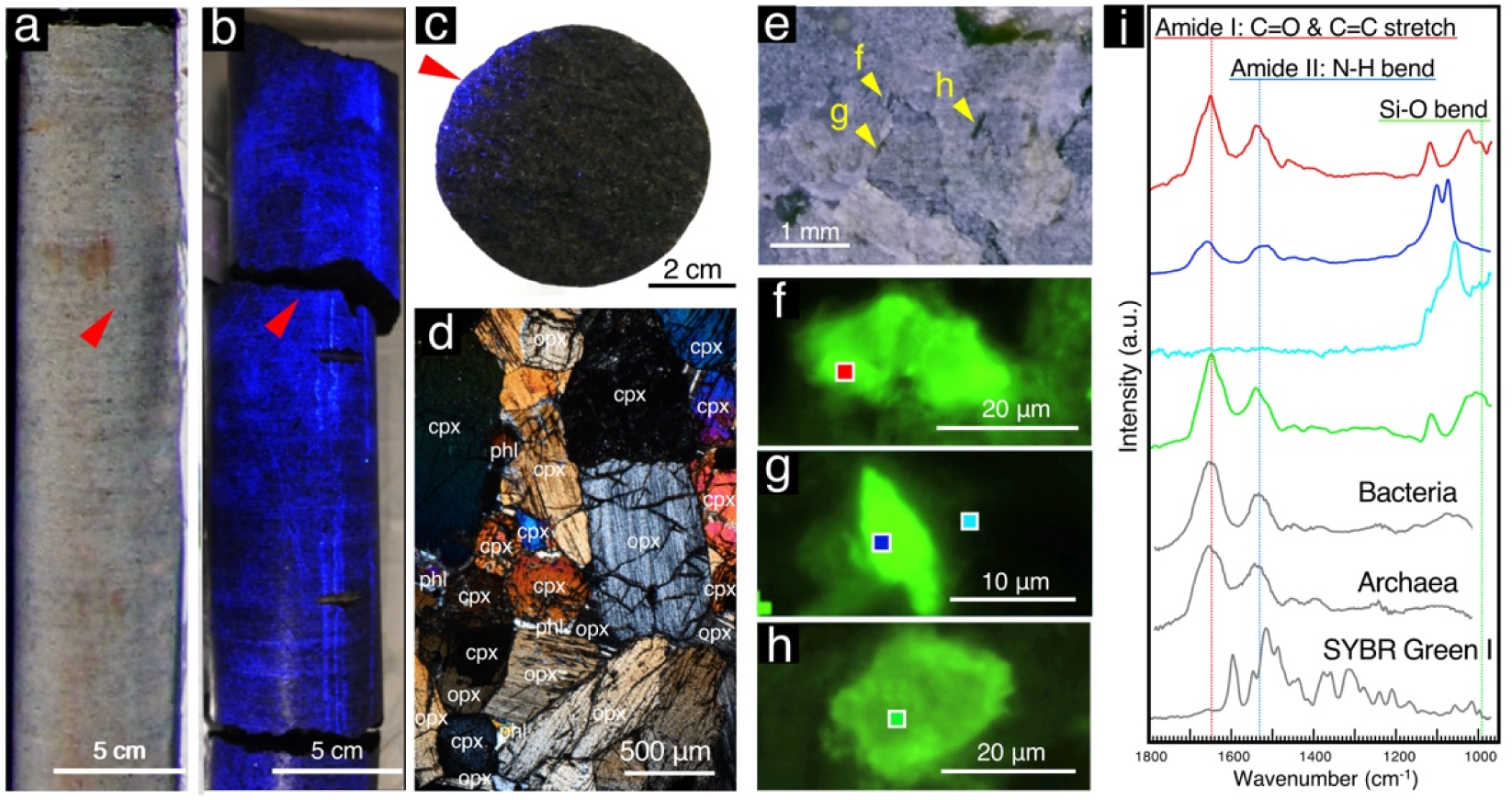
Rock description, contamination discrimination, and in situ microbial detection. **a, b**, Core sample under visible (**a**) and UV light (**b**), showing a crack (red arrowhead) made with a sterile hammer. Blue fluorescence indicates contamination on the edge. **c**, Broken surface under UV light showing contamination where blue fluorescence is visible only at the core edge (red arrowhead). **d**, Thin section (crossed polarizers) displaying orthopyroxene (opx), clinopyroxene (cpx), phlogopite (phl), and quartz. **e**, Prepared rock section with analytical points (yellow arrows). **f–h**, Fluorescence microscopy (SYBR Green I) of green regions, with analytical spots marked. **i**, O-PTIR spectra of the rock (colored squares) compared to control microorganisms (*E. coli*, *N. aerobiophila*, and *M. sedula*).

**Extended Data Fig. 1.**
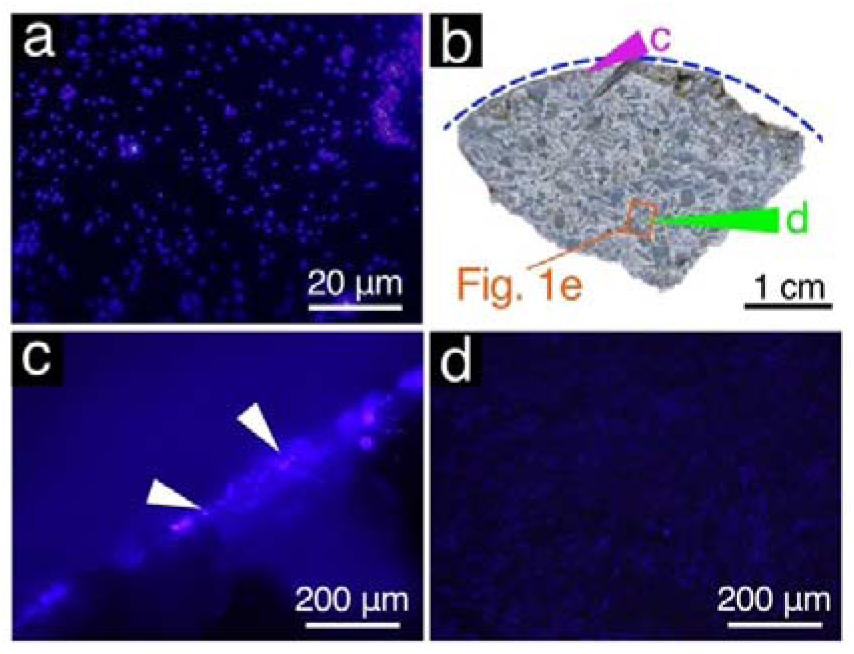
Assessment of drilling fluid contamination using fluorescent microspheres. **a,** High-magnification fluorescence micrograph of fluorescent microspheres within the drilling fluid. **b,** Photograph of a rock section extending from the outer edge to the center of the drill core, prepared using a precision diamond band saw. The blue dashed line indicates the outer edge of the core sample. Colored arrows indicate the areas observed in **c** (magenta) and **d** (green). The orange rectangle corresponds to the area shown in Fig. 1e. **c, d,** High-magnification fluorescence micrographs of the rock section at the locations indicated in **b**. White arrows denote the presence of fluorescent microspheres.

From the core interior and the core exterior, including the outer core surface, we cut 3 mm thick sections with a precision diamond band saw without lubricants (e.g., water or oil) in a clean fume hood with air circulated through a high efficiency particulate air (HEPA) filter (Fig. 1e; Extended Data Fig. 1b). Although microspheres were observed at the outermost edge of the exterior section (Extended Data Fig. 1c), no microspheres were detected in the interior section by fluorescence microscopy (Extended Data Fig. 1d). The interior section was then stained with SYBR Green I and examined using an optical photothermal infrared (O-PTIR) spectroscope equipped with a fluorescence microscope. Microbial signals, indicated by peaks attributed to amide bonds in proteins at 1,530 and 1,640 cm^−1^, were detected at mineral grain boundaries, which coincided with greenish regions stained with SYBR Green I (Fig. 1f–i). In combination with the absence of fluorescent microspheres (Extended Data Fig. 1d), our results demonstrate the presence of indigenous microbial cells at mineral grain boundaries.

We also observed regions with strong greenish signals at or near the edge of the interior section (Fig. 2a), where individual microbial cells were visible with high-magnification fluorescence microscopy (Figs. 2b,c). To obtain independent evidence of the presence of microbial cells, the thin section was studied using scanning fluorescence X-ray microscopy (SFXM) with a soft X-ray beam line at the NanoTerasu synchrotron radiation facility in Japan. Micro-X-ray fluorescence (μ-XRF) mapping of C, N, P, and S with a ca. 3 μm diameter beam revealed multiple loci with co-enrichments of C, N, P, and S at and near the edge of the section (Figs. 2d,e). In addition, the regions with C-, N-, P-, and S-enrichments (Points 1 and 2 in Figs. 2d,e) and a non-enriched control location (Point 3 in Fig. 2d) were subjected to N *K*-edge X-ray absorption near-edge structure analysis (XANES) using a ca. 5 × 25 μm beam. Since this analysis requires a strong X-ray beam to be irradiated at one location for ca. 20 min, we first evaluated the possibility of beam damage to the reference materials (biological materials and inorganic and organic reagents). We confirmed negligible damage for the first and second measurements (Extended Data Fig. 2). The N *K*-edge XANES spectrum from the C-, N-, P-, and S-enriched loci (Point 1) was very similar to those from cultured bacterial cells (*Escherichia coli*) and markedly different from those of the other reference materials, based on the relative intensities of peaks attributed to the imine group at 398.8 eV, pyridine-type heterocyclic compounds at 399.7 eV, NH_4_^+^ at 400.9 eV, and pyrrole-type heterocyclic compounds at 401.2 eV (Fig. 2e,g)^21,22^. Although the N *K*-edge XANES spectrum from the C- and N-enriched loci (Point 2) was similar to that of cultured bacterial cells, the signal-to-noise ratio was low. This result is consistent with the low enrichment levels of C, N, P, and S. Since the control location showed no detectable X-ray absorption features for N, the XANES spectra cannot be attributed to the rock matrix or instrumental artifacts. Taking these results together, we conclude that the unfractured ultramafic rock is colonized by indigenous microbes in the deep subsurface.

**Fig. 2.**
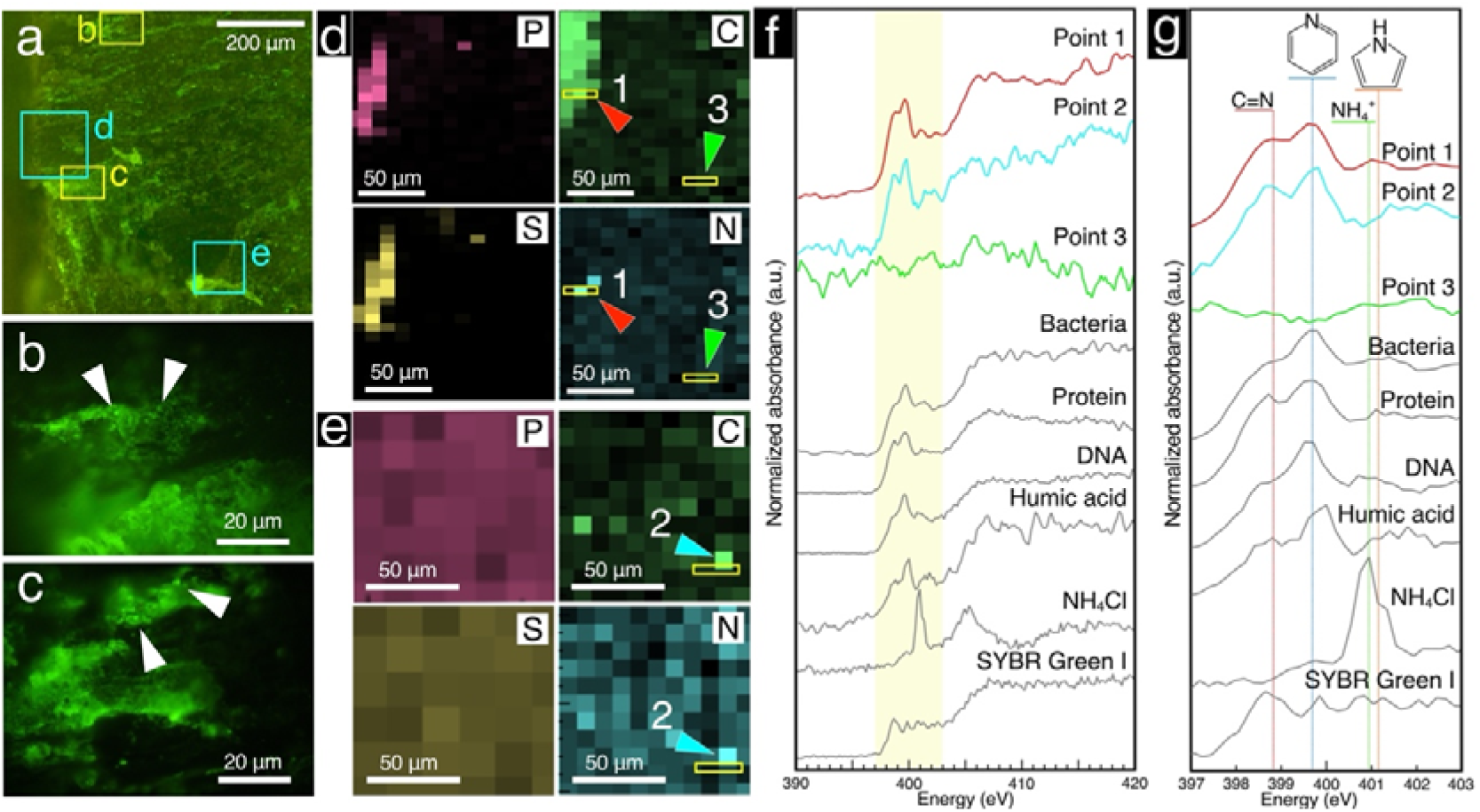
Microbial signals within the rock interior characterized by synchrotron-based soft X-ray spectroscopy. **a**, Fluorescence micrograph of a rock section stained with SYBR Green I. **b, c**, High-magnification fluorescence micrographs showing greenish signals from microbial cells (white arrows); the areas correspond to yellow rectangles in **a**. **d, e**, Elemental maps of C, N, P, and S obtained via micro-X-ray fluorescence (μ-XRF) analysis; the areas correspond to light blue rectangles in **a**. Colored arrows and yellow rectangles indicate locations for X-ray absorption near-edge structure (XANES) analyses. **f**, N *K*-edge XANES spectra of the sample and reference materials. Spectral colors correspond to the arrows in **d** and **e**. Reference materials include cultured bacteria (*Escherichia coli*), protein (albumin), DNA, humic acid, NH_4_Cl, and SYBR Green I. Vertical dotted lines indicate peak positions at 398.8 eV (red), 399.7 eV (blue), 400.9 eV (green), and 401.2 eV (orange). **g**, Magnified N *K*-edge XANES spectra corresponding to the yellow highlighted area in **f**. Absorption peaks represent the imine group at 398.8 eV (red)^21^, pyridine-type heterocyclic compounds at 399.7 eV (blue)^21^, interstitial NH_4_^+^ at 400.9 eV (green)^22^, and pyrrole-type heterocyclic compounds at 401.2 eV (orange)^21^.

**Extended Data Fig. 2.**
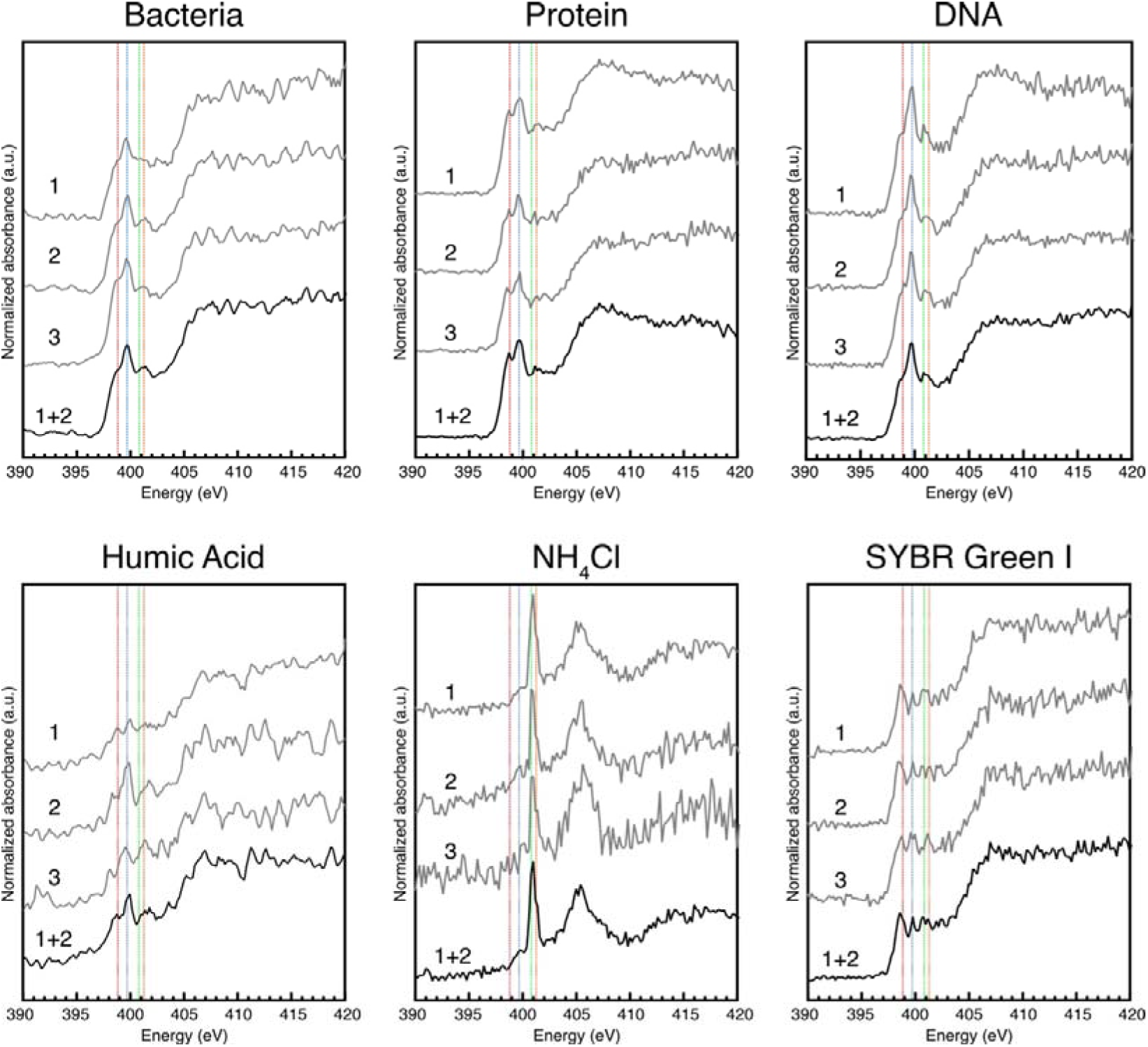
Evaluation of beam damage through repeated N K-edge XANES measurements. N *K*-edge XANES spectra of reference materials (cultured *E. coli*, albumin, DNA, humic acid, NH_4_Cl, and SYBR Green I) from three consecutive scans. Due to negligible beam damage in the initial two scans, they were merged to produce the spectra shown at the bottom and in Fig. 2f. Vertical dotted lines denote peaks at 398.8 (red), 399.7 (blue), 400.9 (green), and 401.2 eV (orange).

### Microbial habitat around pyroxene grain boundaries with phyllosilicate minerals

To characterize the habitat of the detected microbes, we performed mineralogical and geochemical analyses of the regions where the SFXM data demonstrated the presence of microbial colonization. Environmental scanning electron microscopy (ESEM) coupled with energy-dispersive X-ray spectroscopy (EDS) revealed microbial signals located at orthopyroxene grain boundaries in contact with intercumulus grains interpreted to be phlogopite (Mg-rich mica) based on the mineral’s EDS spectra (Extended Data Fig. 3a,b). We also found microbial signals spatially correlated with the rims of the phlogopite grains, where the EDS spectra indicated K depletion (Extended Data Fig. 3c). In addition, a region enriched with iron and sulfur was observed near the phlogopite grains (Extended Data Fig. 4a,b). We performed S *K-*edge XANES analysis for the Fe- and S-enriched region, and the spectra obtained from this region were identical to those of pyrrhotite (iron monosulfide) (Extended Data Fig. 4c).

**Extended Data Fig. 3.**
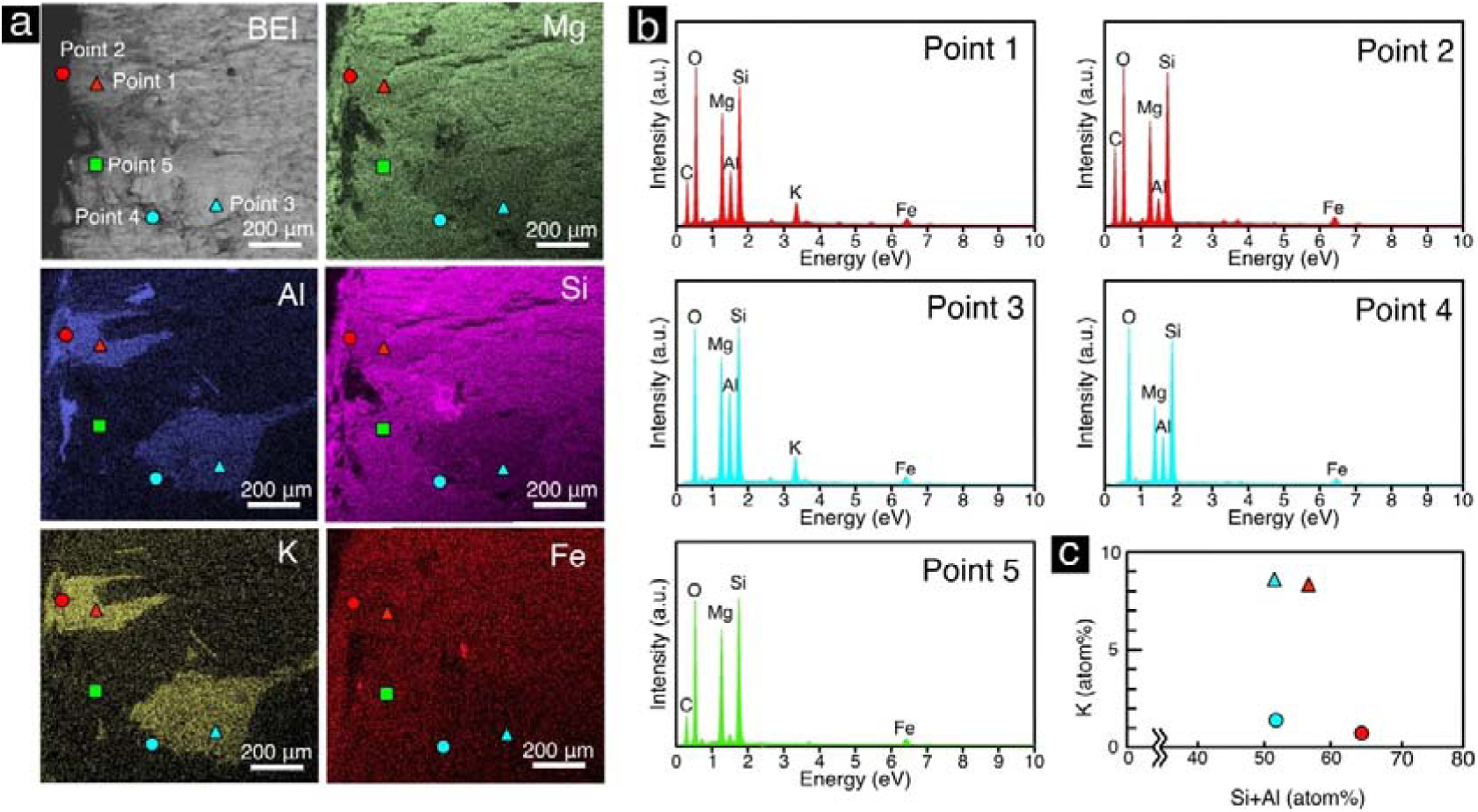
Mineralogical characterization of microbially colonized regions. **a**, Back-scattered electron (BSE) image and elemental maps (Mg, Al, Si, K, and Fe) obtained via environmental scanning electron microscopy with energy-dispersive X-ray spectroscopy (ESEM-EDS). The mapped area corresponds to Fig. 2a and Fig. 3a,b. Symbols indicate analytical locations. **b**, Representative ESEM-EDS spectra for points 1–5 in **a**. **c**, Binary plot of K versus Si+Al (atomic %) showing K-depletion along the rims of phlogopite-like grains. Plot symbols correspond to those in **a**.

**Extended Data Fig. 4.**
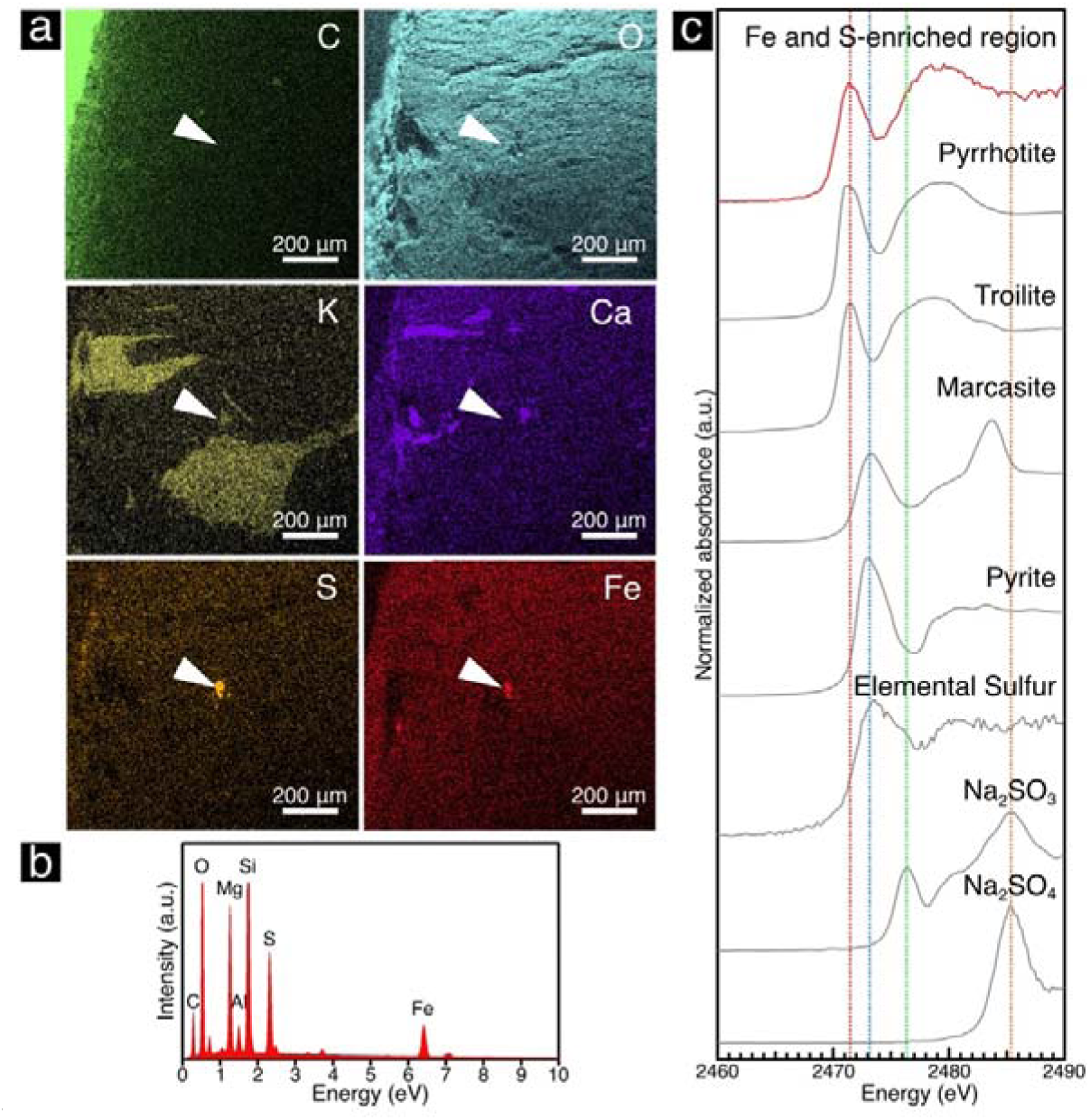
Characterization of an iron- and sulfur-enriched region near phlogopite grains. **a,** Element maps of C, O, K, Ca, S, and Fe acquired via environmental scanning electron microscopy with energy dispersive X-ray spectroscopy (ESEM-EDS). White arrows indicate the location of the iron- and sulfur-enriched region. The maps cover the same area shown in Fig. 3a,b and Extended Data Fig. 3a. **b,** EDS spectra extracted from the iron- and sulfur-enriched region. **c,** S *K*-edge XANES spectra of the iron- and sulfur-bearing grain and reference materials (pyrrhotite, troilite, marcasite, pyrite, elemental sulfur, Na_2_SO_3_, and Na_2_SO_4_). Vertical dotted lines mark peak positions at 2471.4 eV (red), 2473.2 eV (blue), 2476.3 eV (green), and 2485.3 eV (orange).

We also employed powder X-ray diffraction (XRD) analysis of the whole rock to identify the mineralogy of the sample. In an XRD pattern of the whole rock, peaks were attributed to orthopyroxene, clinopyroxene, quartz, and phlogopite (Extended Data Fig. 5a). An XRD pattern of the clay-sized fraction of the sample showed peaks at 10.0 Å and 14.3 Å (Extended Data Fig. 5b). After treatment with ethylene glycol, no peak shift was observed for the peaks at 10.0 Å and 14.3 Å, indicating the absence of smectite-group minerals^23^. As the 14.3 Å peak was shifted to 10.0 Å after heat treatment at 500°C, the 14.3 Å peak was identified as a vermiculite-group mineral^23^.

**Extended Data Fig. 5.**
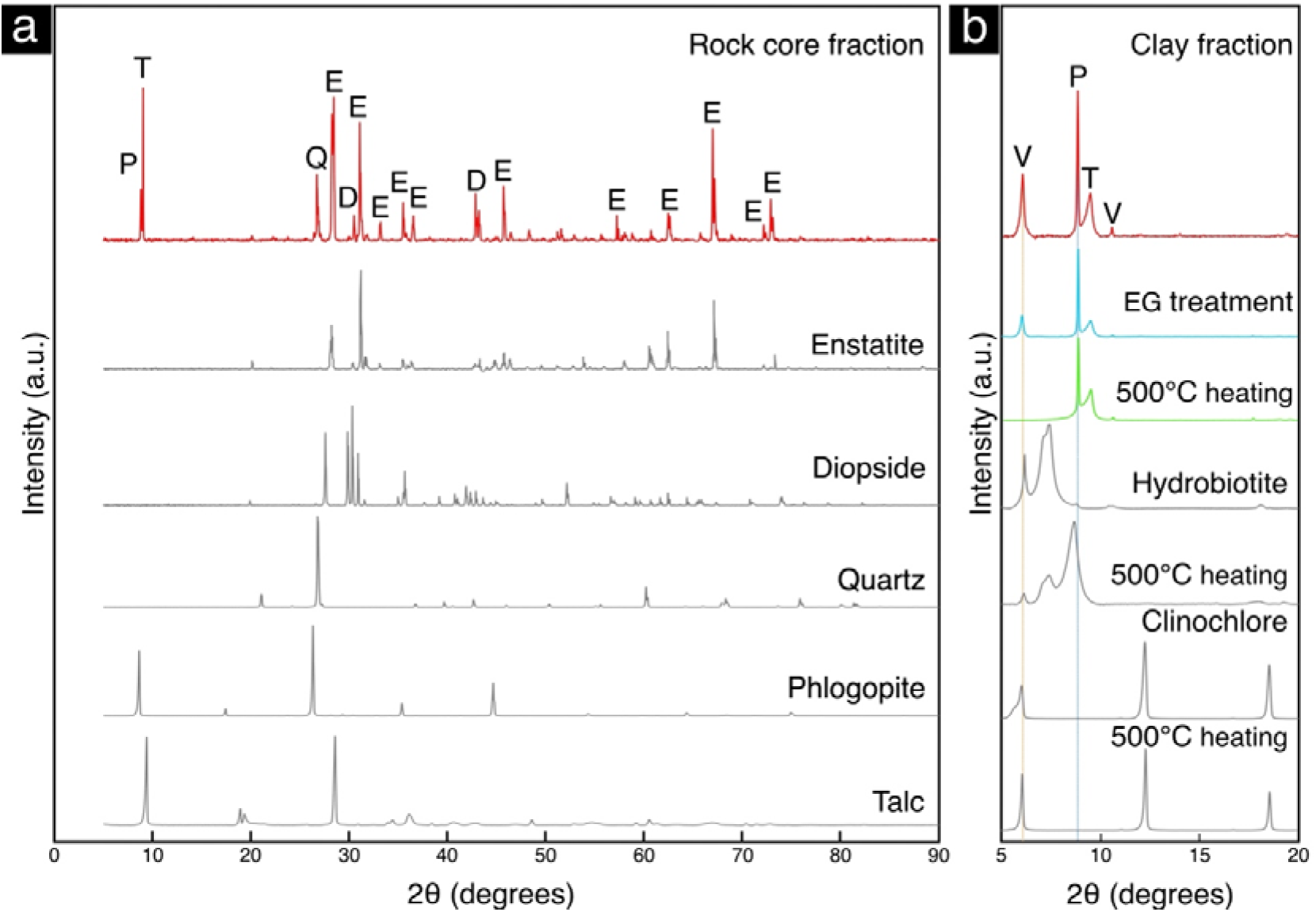
Mineral identification by X-ray diffraction analysis. **a,** Powder X-ray diffraction (XRD) patterns of the core sample and reference minerals. Peak labels are as follows: E, enstatite; D, diopside; Q, quartz; P, phlogopite; T, talc; and V, vermiculite. **b,** Powder XRD patterns of the oriented clay-size fraction from the core sample before and after ethylene glycol (EG) treatment and heating (500 °C for 1 hour). Hydrobiotite (an interstratification of Fe-rich mica and vermiculite)^66^ and clinochlore (Mg-rich chlorite) are shown as references to demonstrate the diagnostic collapse of the vermiculite interlayer spacing upon heating. Vertical lines indicate the peak positions at 10.0 Å (blue) and 14.3 Å (orange)

To further constrain the identity of the phlogopite grains and the K-depleted areas along their rims, which had a spatial association with microbial signals, we performed in situ mineralogical analysis using SFXM with a soft X-ray at the SPring-8 synchrotron radiation facility in Japan. μ-XRF mapping of Al with a ca. 2-μm diameter beam (Fig. 3b) confirmed the locations where previous SFXM and ESEM-EDS analyses indicated the presence of microbial cells (Fig. 2) in association with the rims (Extended Data Fig. 3). Al *K*-edge XANES spectra are known to be distinct among rock-forming minerals^24^, but there has been no systematic study to compare Al *K*-edge XANES spectra from various phyllosilicate minerals. In this study, we found that Al *K*-edge XANES are also sensitive to different phyllosilicate minerals (Fig. 3c). The Al *K*-edge XANES spectra of the phlogopite grains were similar to that of phlogopite (Fig. 3c). In addition, spectra identical to that of hydrobiotite, a mineral consisting of interstratified vermiculite and biotite (Fe-rich mica), were obtained from the K-depleted rims of the phlogopite grains with microbial signals (Fig. 3c). As vermiculite is generally transformed from phlogopite by loss of interlayer K by hydrothermal alteration and/or weathering^9^, it is reasonable to expect to obtain microbial signals from the K-depleted rims of the phlogopite grains. The presence of vermiculite in the sample is supported by the presence of vermiculite and phlogopite peaks in the powder XRD pattern of the sample (Extended Data Fig. 5).

**Fig. 3.**
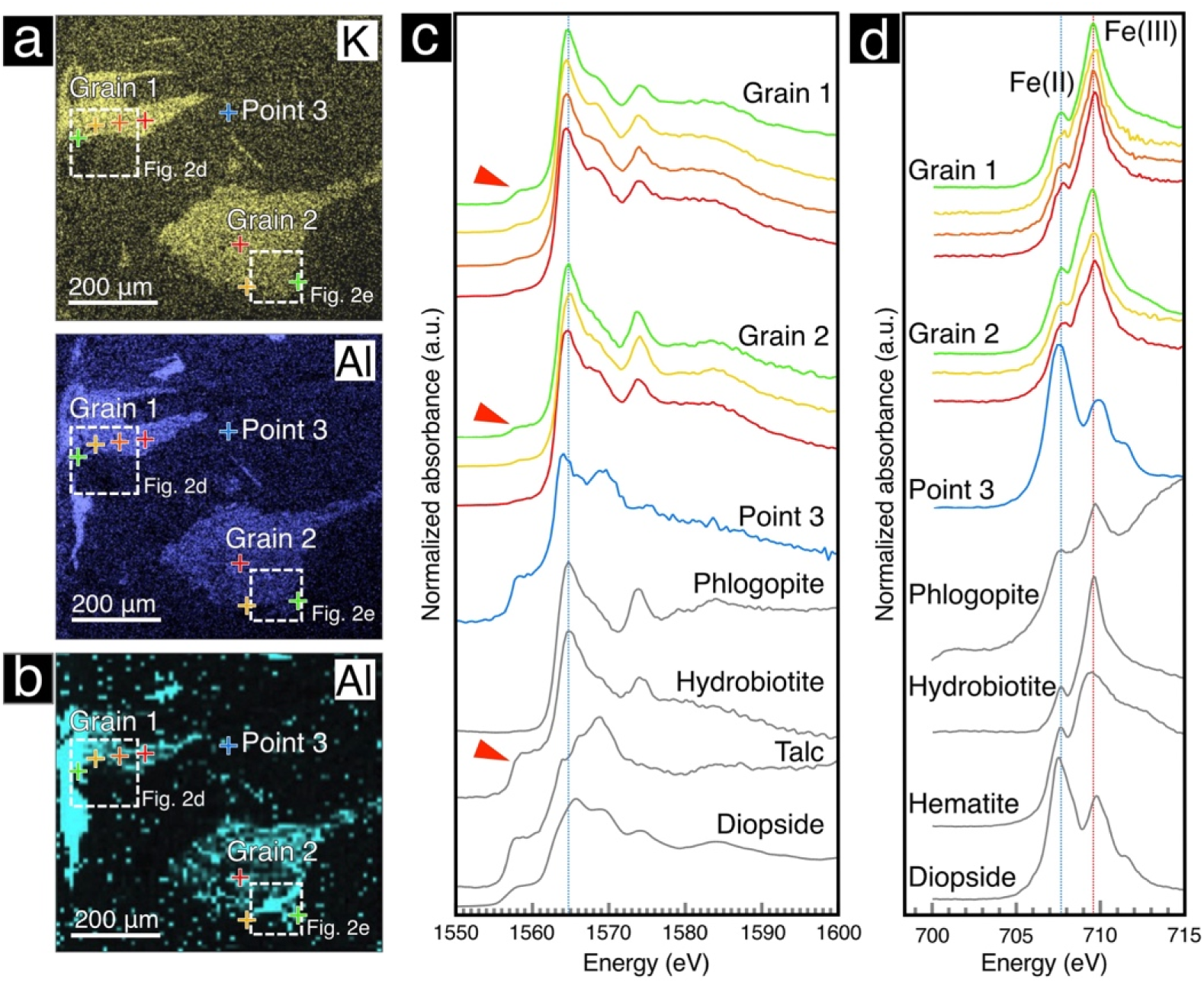
Mineralogical and geochemical features associated with microbial colonization. **a,** Element maps of Al and K obtained by environmental scanning electron microscopy with energy dispersive X-ray spectroscopy (ESEM-EDS). **b,** Synchrotron-based micro-X-ray fluorescence (μ-XRF) map of Al for the same region shown in **a**. The mapped area corresponds to Fig. 2a. **c, d,** X-ray absorption near-edge structure (XANES) spectra obtained from the rock section and reference minerals at the Al *K*-edge (**c**) and Fe *L_3_*-edge (**d**). Pre-edge structures (red arrows in **c**) indicate the presence of vermiculite in hydrobiotite. Spectral colors correspond to the analyzed points in **a** and **b**. Reference minerals include phlogopite, hydrobiotite, talc, saponite, nontronite, montmorillonite, hematite, and diopside.

Fe *L_3_*-edge XANES analysis was performed on the same locations as the Al *K*-edge XANES analysis to distinguish the valence state of iron based on the spectra at ca. 707 and ca. 710 eV^25^. The relative peak intensities obtained indicated that Fe(III) was abundant at the phlogopite grains, whereas Fe(II) dominated at the adjacent pyroxenite grains (Fig. 3d). We also performed Fe *L_3_*-edge XANES analysis from the interior to the exterior of the phlogopite grains. The relative peak intensities indicated the relative abundance of Fe(II) at the phlogopite rims (Fig. 3d), which is consistent with the microbial reduction of Fe(III) mediated at the phlogopite rims associated with microbial signals (Fig. 2).

As there was the possibility that Fe(II) in the phlogopite grains and their vermiculite-bearing rims was oxidized by air exposure during sample preparation and subsequent analyses^26^, we prepared a new 3 mm thick rock section using a precision diamond wire saw placed inside an Ar-purged glove box (Fig. 4a). Fe *K*-edge XANES analysis of the rock section in an He-purged sample holder was performed using SFXM with hard X-rays at SPring-8. μ-XRF mapping of K revealed the presence of phlogopite-like grains (Fig. 4b). The valence state of iron around the phlogopite-like grains was determined based on the Fe *K*-edge position^27^. The Fe *K*-edge position was shifted toward that of an Fe(III) standard^28^ from the exterior to the interior of the phlogopite grain (Fig. 4c,d,e). The mineral identity was confirmed by Al *K*-edge XANES analysis with soft X-rays (Fig. 4f). This result is consistent with the shifts in the relative peak intensities of Fe(II) and Fe(III) in the Fe *L_3_*-edge XANES spectra from the air-exposed rock section (Fig. 3d). Given that the oxidation of iron in biotite in air typically proceeds at temperatures above 400°C^30^, it is unlikely that the presence of Fe(III) in the phlogopite grains and their vermiculite-bearing rims was an artifact introduced after core processing and subsequent characterizations of the rock sections.

**Fig. 4.**
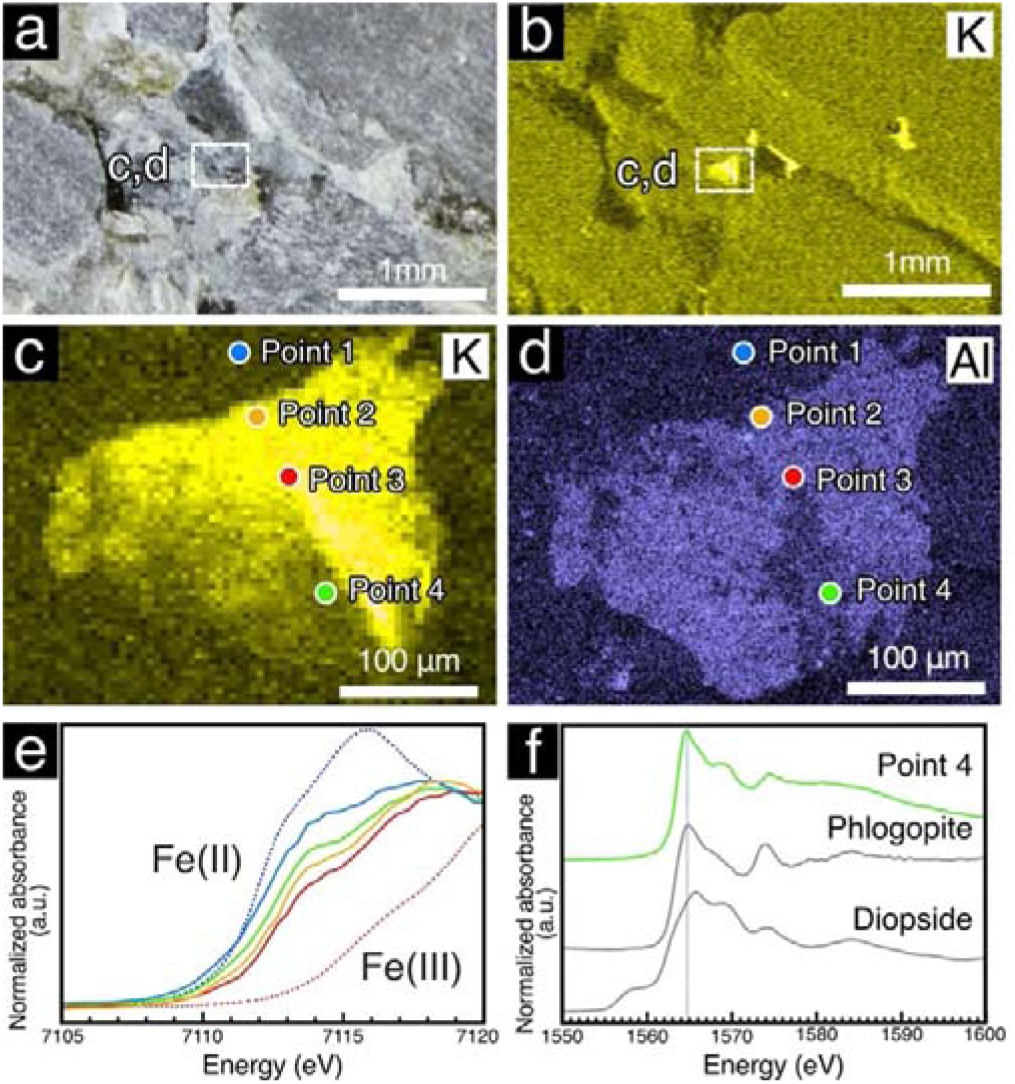
Internal variation in the iron valence state inside and outside a phlogopite grain determined after minimal exposure to air. **a**, Photograph of the rock section prepared by a diamond wire saw in an Ar-purged glove box. **b**, Element map of K obtained by synchrotron-based micro-X-ray fluorescence (μ-XRF) analysis. The area corresponds to the rectangle in **a**. **c**, Higher magnification map of K obtained by the same μ-XRF analysis. The area corresponds to the rectangle in **a** and **b**. **d**, Element map of Al obtained using environmental scanning electron microscopy with energy dispersive X-ray spectroscopy (ESEM-EDS). The area corresponds to the rectangle in **a** and **b**. **e**, Fe *K*-edge X-ray absorption near-edge structure (XANES) spectra obtained inside and outside a K- and Al-enriched grain in the rock section and references. The spectral colors correspond to points indicated in **c** and **d**. FeCl_2_^29^ and nontronite^28^ were used as the references for Fe(II) (blue dotted line) and Fe(III) (red dotted line), respectively. **f**, Al *K*-edge XANES spectra obtained from the K- and Al-enriched grain and reference materials (phlogopite and diopside).

## Discussion

### Magmatic-to-aqueous evolution forming microbial habitats

The presence of microbial life at the grain boundaries of hydrous-altered phlogopite suggests that the suitability of the habitat depends on the interaction of magmatic minerals and fluids. Our results suggest the following possible scenario, as illustrated in Fig. 5.

**Fig. 5:**
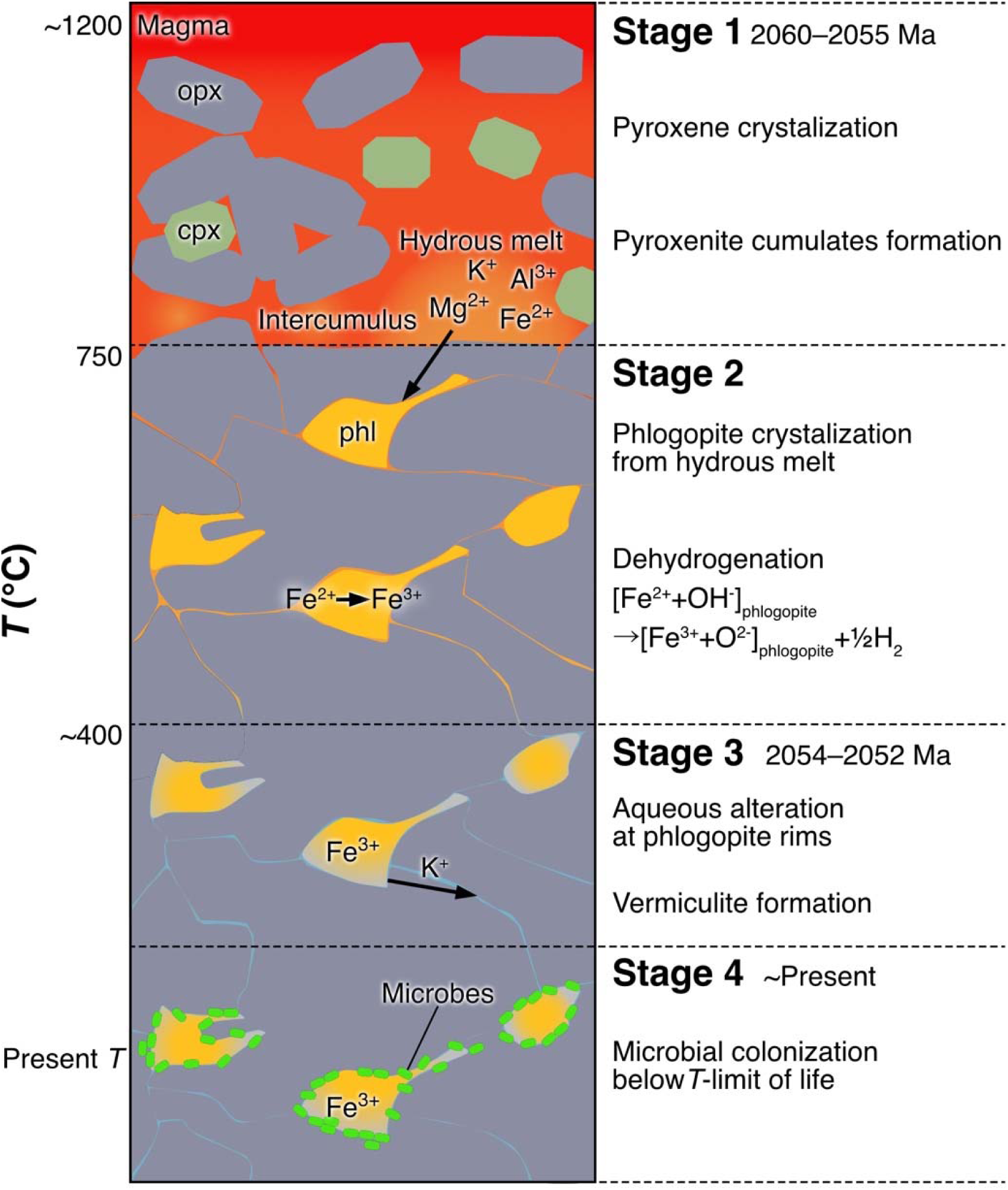
Conceptual model for the formation of deep microbial habitats in pyroxenite of the Bushveld Igneous Complex. The model comprises four stages. **Stage 1**: Orthopyroxene (opx, grey) and clinopyroxene (cpx, green) crystallize, forming a cumulate layer with interstitial residual melt enriched in water and incompatible elements (e.g., K). **Stage 2**: Phlogopite (phl, yellow) crystallizes from residual, hydrous melt in the intercumulus space and undergoes dehydrogenation upon cooling. **Stage 3**: The rims of phlogopite react with aqueous fluid, forming vermiculite while retaining Fe(III). **Stage 4**: Following cooling to temperatures below the limit of life, microbes colonize the altered phlogopite rims, with energy provided by the reduction of Fe(III).

Stage 1: Crystallization and accumulation of primary pyroxene crystals from magma (e.g., <ca. 1,200°C) ^31^.

Stage 2: Crystallization of intercumulus phlogopite from a hydrous interstitial melt (<750°C) ^32^.

Stage 3: Transformation of phlogopite to vermiculite by hydrous alteration below the critical point of water in saline fluid (ca. 400°C)^33^. Based on the cooling history of the intrusion, this temperature was reached about 10 million years after the intrusion occurred^34^.

Stage 4: Further cooling to the present temperature. The rock sample studied is spatially associated with brackish groundwater inflowing into the drilled borehole through fractures at various depths. The temperature of the brackish groundwater was measured to be ca. 37°C in the borehole^19^.

Between Stages 2 and 3, cooling of the phlogopite is expected to have driven the dehydrogenation reaction, in which Fe(II) oxidation in the crystal lattice and H_2_ generation proceed at 640°C–750°C^8^:

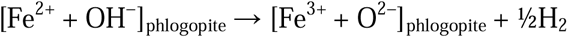

Fe(III) in the phlogopite structure is subsequently maintained, even after the phlogopite is transformed into vermiculite^35^. This scenario is supported by the presence of Fe(III) throughout the phlogopite grains (Fig. 4c–f), because Fe(II) oxidation by O_2_-bearing fluid tends to oxidize the grain rim. This aqueous process is unlikely to have occurred, given the low permeability of the unfractured host rock^19^. Additional evidence against fluid-driven oxidation is the preservation of redox-sensitive iron monosulfide (pyrrhotite) found near the phlogopite grains (Extended Data Fig. 4). Given that pyrrhotite is rapidly oxidized and dissolved by O_2_ in fluid^36^, it is unlikely that the rock was exposed to O_2_-bearing fluid during Stage 4.

Since abiotically generated microbial energy sources such as H_2_, CH_4_, and organic acids are commonly present as microbial energy sources in the deep subsurface^37^, microbes can survive by oxidizing these energy sources using the structural Fe(III) in phlogopite and/or vermiculite as oxidants. In the case of the oxidation of H_2_ coupled to the reduction of Fe(III) in vermiculite, H_2_O is produced as a waste product. This metabolic pathway is commonly linked with chemolithoautotrophic growth, during which CO_2_ is taken up by microbial cells. Pyroxenite lacks fractures or veins that would allow fluid transport of metabolic nutrients and the removal of waste^3^. Given that the rims of phlogopite grains appear to serve as a low-permeability habitat, microbial survival requires the prevention of pore occlusion by metabolic byproducts in addition to the redox gradients exploited by microbial metabolisms^3^. As Fe is retained within the crystal lattice of vermiculite after the reduction of Fe(III), pore occlusion by Fe(III) reduction is unlikely^38^. Given that H_2_ is available at grain boundaries without needing to be transported by fluid flow, it is reasonable to find microbial colonization at the vermiculite-bearing rim in association with Fe(III) in solid rock without fractures or veins.

Although determining the precise timing of microbial colonization remains challenging, the lack of metamorphic overprinting and the absence of fracture-driven fluid flow suggest that the internal redox gradient was established after the intrusion cooled 2.05 billion years ago (Stage 3). The preservation of the internal redox gradient with no equilibration by any subsequent fluid flow provides a geochemical constraint that limits the potential for recent biological ingress and drilling-fluid contamination. The longevity of this isolated habitat raises questions regarding the energetic limits of life. We propose that Bushveld ultramafic rock provides a continuous, albeit minimal, energy flux for cell maintenance, rather than cell growth from the reduction of Fe(III)-bearing phyllosilicate minerals^39,40^.

### Analytical advancement in biosignature detection from ultramafic rocks

Scientific drilling in the search for deep microbial life has repeatedly targeted ultramafic rocks in the oceanic and continental crust^41^ because aqueous alteration of ultramafic rocks provides not only chemical energy but also mineral catalysts for prebiotic synthesis and the emergence of life^42^ and for living and fossil rock-hosted microbes^3^. A number of techniques have been developed to detect biosignatures. Raman spectroscopy has been applied to hydrogarnet grains in ca. 1-million-year-old (Ma) seafloor ultramafic rocks from the Mid-Atlantic Ridge^43^ and brucite veins in 120 Ma subseafloor ultramafic rocks at the Iberian Margin^44^. Despite the presence of sharp peaks attributed to aliphatic compounds and functional groups, such as amides, usually associated with biopolymers, such as proteins and lipids, Raman biosignatures from ultramafic rocks differ from those of cultured microbial cells. These differences are attributed to the type and metabolic activity of the cells, along with preservation with aging^43^. Raman spectra from microbial filaments morphologically preserved in calcium carbonate veins in ultramafic rocks in the ca. 100 Ma Samail Ophiolite, Oman^45^, and in ca. 1 Ma seafloor ultramafic rocks from the Mid-Atlantic Ridge^46^, lack sharp peaks from biopolymers. Furthermore, they show broad peaks attributed to organic material without structural information, which could reflect the poor preservation potential of biopolymers in calcium carbonate minerals and/or interference by autofluorescence from calcium carbonate minerals. Autofluorescence interference is also a problem affecting the use of Raman spectroscopy to obtain organic signals from microbial cells associated with phyllosilicate minerals^47^.

Fourier transform infrared (FT-IR) spectroscopy has been widely used to detect organic molecules associated with phyllosilicate minerals in ultramafic rocks. In ultramafic rocks from the ca. 2 Ma Atlantic Massif, an Mg-rich phyllosilicate mineral, saponite, is associated with abiotic amino acid synthesis, as characterized by synchrotron-based FT-IR spectroscopy with a 5 × 5 μm aperture^42^. In our study, spectra comparable to those from FT-IR were obtained from the Bushveld rock using O-PTIR spectroscopy. In this technique, thermal expansion by infrared illumination is detected by a green laser (532 nm) with a beam diameter of ca. 0.5 μm^48^. This O-PTIR spectroscopy can achieve a high spatial resolution equivalent to Raman spectroscopy without autofluorescence interference (Fig. 1i). The O-PTIR spectra obtained from the pyroxene grain boundaries in this study are similar to those of cultured microbial cells (Fig. 1i) and distinct from abiotic amino acids^49^. Given that phyllosilicate minerals preserve organic molecules from Archean siliciclastic microbial mats^50^, the possibility that O-PTIR spectroscopy detected fossil microbial cells that were exceptionally well preserved by coexisting phyllosilicate minerals is not excluded but is unlikely.

We further advanced analytical techniques by applying SFXM to validate the biogenic origin of fluorescence microscopic and O-PTIR spectroscopy signals in the deep ultramafic rock, as well as the phyllosilicate mineral identity estimated by bulk XRD and ESEM analyses. In our previous SFXM-based study^51^, we revealed N-bearing organic matter in spatial association with the reduction of Mn(IV) and Ce(IV) at a deep-sea ferromanganese crust surface. This organic matter is characterized by the enrichment of aromatic and carboxylate groups in N *K-*edge XANES spectra, which are clearly distinct from microbial cells. To our knowledge, the present study is the first demonstration that SFXM can be used to detect the biosignatures of rock-hosted microbes. A similar synchrotron-based method called scanning transmission X-ray microscopy (STXM) is commonly used to detect microbial cells based on N *K*-edge XANES spectra^52,53^. STXM detects X-ray absorption after an incident beam is transmitted through a sample, whereas in SFXM, the X-ray absorption is detected using fluorescence X-rays emitted from a sample without transmission^54^. An STXM sample needs to be thinned to ca. 100 nm, which can be problematic for void spaces around mineral grains loosely filled with microbe–mineral assemblages. In contrast, an SFXM sample only requires a flat surface prepared by cutting with a precision diamond band and/or wire saw, which enables the preservation of fragile rock habitats.

Our synchrotron-based analysis is also critical to support the indigenous nature of the observed microbial colonization. N *K*-edge XANES spectra identical to microbial cells (Fig. 2) were obtained at the mineral grain boundaries, where the aqueous alteration and the internal redox gradient essential to microbial survival in an unfractured rock matrix were reliably localized by μ-XRF analysis and Al and Fe *K*-edge XANES analyses (Figs. 3 and 4). The spatial congruency, in combination with our rigorous contamination-control protocols using fluorescent microspheres, strongly argues against drilling-induced contamination (Fig. 1b,c and Extended Data Fig. 1).

### Implications for the search for rock-hosted life on Mars

Our findings demonstrate that the ancient, unmetamorphosed ultramafic rock in the Kaapvaal Craton hosts an isolated microbial habitat. This discovery challenges the conventional view of the deep-rock biosphere in which microbial life is strictly dependent on nutrient delivery through fracture networks^3,36^. Instead, the self-sustaining nature of this habitat—driven by the intrinsic redox potential of Fe-bearing phyllosilicate minerals—suggests that life can persist in a state of extreme isolation for as long as mineral–water reactions continue to yield chemical energy. Such a mechanism has profound implications for the search for life on Mars.

Due to a lack of plate tectonics, metamorphic overprinting is very limited in Martian rocks. Based on the similarity to the Bushveld Igneous Complex, the Noachian (3.7–4.1 Ga) Columbia Hills on Mars have been regarded as a layered intrusion^10^. This formation was exhumed by a meteorite impact that formed the Gusev Crater in the Hesperian (3.7–3.9 Ga) and underwent aqueous alteration^55^. The presence of olivine- and pyroxene-rich cumulates has also been reported in the Jezero Crater^56^, with varying age estimates from 1.4–3.45 Ga^57^. Considering these time periods, our results from the Bushveld Igneous Complex suggest that the Martian ultramafic rocks may still harbor ancient microbial lineages or their detectable biosignatures. The Perseverance rover, equipped with a deep-UV Raman spectrometer with a beam diameter of 350 μm, detected one of the broad Raman bands attributed to C=C stretching vibrations at ca. 1,600 cm^−1^ from the pyroxene-rich cumulate from the Jezero Crater^58^. In the Raman spectra with the broad band at ca. 1,600 cm^−1^, a peak at ca. 1,080 cm^−1^ was also present and was provisionally assigned to silicate, due to an inability to resolve silicate phases^59^. For future missions to search for Martian life, the use of O-PTIR spectroscopy would enable better characterization of organic molecules in aqueously altered ultramafic rocks, as demonstrated in our application on Earth, which was confirmed by synchrotron-based spectroscopy.

### Microbial survival with minimal evolution in ultramafic intrusions older than 2 Ga

This study provides a new geological template for searching for ancient microbes from long-term habitats on Earth. Archean cratons other than those previously mentioned also host unmetamorphosed ultramafic layered intrusions, such as the Munni Munni Complex (ca. 2.93 Ga) in the Pilbara Craton in Australia^60,61^, the Great Dyke (ca. 2.58 Ga) in the Zimbabwe Craton in Zimbabwe^62,63^, and the Näränkävaara layered igneous complex (ca. 2.44 Ga) in the Karelian Craton in Finland^64^. In all of these ultramafic intrusions, the formation of intercumulus phlogopite is evident. Our study strongly suggests that deep microbial life that depends on the aqueous alteration of Fe(III)-bearing phlogopite may be ubiquitous in unmetamorphosed ultramafic layered intrusions formed in the Neoarchean to Paleoproterozoic eras. In addition to crustal fluid isolated for 1.2 billion years, the Kaapvaal Craton harbors a subsurface bacterial species that has undergone minimal evolution for 55–165 million years in fractured metamorphic rocks^65^. Our Bushveld research targets microbial genomes that potentially preserve evolutionarily primitive features for billions of years in the geologically and tectonically stable subsurface environment.

## Methods

### Drilling and on-site core handling procedures

A rotary core barrel with a diamond bit was used for drilling. The drilling fluid was composed of locally sourced water and a high-molecular-weight polymer viscosifier (AMC CAP 21^TM^, AMC/IMDEX Ltd., Balcatta, WA, Australia) to reduce friction and improve core recovery. Samples for microbiological study were collected every 100 meters during drilling, or approximately once every 10 days. Before the planned sampling, fluorescent microspheres (Invisible Blue, DayGlo Color Corp., pigment SPL-594NXC, 0.25–0.45 μm) were added to the drilling fluid in a tank. The analyzed sample from ca. 814 m was drilled and collected on August 5th, 2024. The core sample (Sample ID: 5067_3_B_236R_2_WR:51-87, IGSN ICDP5067EXFA001) was rinsed with deionized water three times shortly after reaching the surface and then lightly flamed with a gas torch, following procedures developed in previous studies^6,7^. Blue fluorescence from fluorescent microspheres on the core surface was observed with a 365 nm-UV light (Nichia Corp., Tokushima, Japan). The decontaminated core sample was stored at −18°C in an Al-lined plastic bag filled with N_2_ gas. Drilling fluid was collected from the mud tank and stored at −18°C.

### Microscopic enumeration of fluorescence microspheres and microbial cells

The core sample was cracked with a flame-sterilized hammer, and the intrusion of fluorescent microspheres into the core interior was visualized by observation of the freshly fractured cross-section. Fluorescent microspheres in the drilling fluid were collected on a 13-mm-diameter polycarbonate filter (0.2-μm pore size; Merck Millipore, Darmstadt, Germany) and counted in triplicate (*n* = 3) using a fluorescence microscope (Olympus BX51, Tokyo, Japan) equipped with a CCD camera (Olympus DP71) and image processing software (Lumina Vision, Mitani Shoji, Tokyo, Japan). Microbial cells in the same drilling fluid were stained with SYBR Green I (Takara-Bio, Inc., Shiga, Japan) and counted (*n* = 3). For rock analysis, the interior and exterior portions were aseptically separated using a sterilized rock trimmer (Iwamoto Mineral Co., Ltd., Tokyo, Japan) and ground into powder with a sterilized titanium-cylinder mortar. Fluorescence microspheres were extracted by suspending 0.5 cm³ of the powder in 2 ml of deionized water, followed by 30 s of ultrasonication. A 0.05-ml aliquot of the suspension was diluted to 2 ml with deionized water, passed through a 13-mm-diameter polycarbonate filter (0.2-μm pore size), and counted using the fluorescence microscope system (*n* = 3). The detection limit, defined as the mean ± three standard deviations (SD) of five replicate blanks (*n* = 5), was 0.75 ± 1.3 × 10^3^ microspheres cm^−3^. The rock interior was cut into a 3-mm-thick section using a precision diamond band saw (DWS 3500P; Meiwa Fosis Corp., Tokyo, Japan) in a clean booth flushed with HEPA-filtered air and observed via fluorescence microscopy.

### Detection of microbial cells in the rock core interior

The 3-mm-thick rock section was stained with SYBR Green I and examined for microbial cells using a fluorescence microscope system. Optical-photothermal infrared (O-PTIR) spectroscopy equipped with fluorescence microscopy (mIRage-LS, Photothermal Spectroscopy Corp., Santa Barbara, USA) was used to detect greenish fluorescent signals from microbial cells stained with SYBR Green I, from which O-PTIR spectra diagnostic of microbial cells were automatically obtained using the feature finder function. A continuous-wave 532-nm laser was used as a probe beam with submicron spatial resolution, whereas a tunable quantum cascade laser (950–1800 cm^−1^; 2 cm^−1^ spectral resolution; 10 scans per spectrum) served as the pump beam to obtain the mid-IR spectra. O-PTIR spectra were also obtained from reference materials such as cultured cells of *Escherichia coli* (*E. coli*; NBRC13168), co-cultured cells of *Nanobdella aerobiophila* (*N. aerobiophila*) strain MJ1, *Metallosphaera sedula* (*M. sedula*) strain MJ1HA (JCM33617), and SYBR Green I.

Micro-X-ray fluorescence (μ-XRF) mapping was performed using a scanning fluorescence X-ray microscope (SFXM) at the BL13U beamline of NanoTerasu. The incident soft X-ray beam was focused to a diameter of ca. 3 μm using a Wolter mirror. The energy and intensity of the fluorescent soft X-rays emitted from the sample were measured using a silicon drift detector in partial fluorescence yield (PFY) mode. Excitation energies of 3000 eV and 700 eV were applied for the simultaneous mapping of P/S and C/N, respectively. Subsequently, N *K*-edge X-ray absorption near-edge structure (XANES) spectra were acquired from the rock section in the 390–420 eV range with a 0.2 eV step. Reference materials, including *E. coli* cells, bovine serum albumin (fatty acid-free, FUJIFILM Wako), salmon sperm DNA (FUJIFILM Wako), International Humic Substance Society (IHSS) Nordic Aquatic Humic Acid (1R105H), SYBR Green I, and ammonium chloride (NH_4_Cl, FUJIFILM Wako), were similarly measured. The beam size for N *K*-edge XANES was ca. 5 μm (vertical) × 25 μm (horizontal). To evaluate beam-induced damage, N *K*-edge XANES spectra were acquired three times at the same spot on the reference materials.

### Petrologic, mineralogical, and geochemical characterizations

A petrographic thin section with a thickness of ca. 30 μm was prepared from the rock core sample after embedding in LR White resin (London Resin Co. Ltd., Aldermaston, England) and then examined under a polarized microscope (BX51-P; Olympus Co. Ltd., Tokyo, Japan). For X-ray diffraction (XRD) analysis, the rock core sample was pulverized using a tungsten carbide mortar and pestle. The clay-sized fraction was isolated by dispersing the powder in deionized water, followed by centrifugation at 3000 rpm for 5 min and freeze-drying of the supernatant. Both whole-rock and clay-fraction samples, along with reference minerals, were analyzed using a RINT-2100 X-ray diffractometer (Rigaku Co. Ltd., Tokyo, Japan) operated at 40 kV and 30 mA with Cu Kα radiation (λ = 1.5406 Å). For randomly oriented samples, XRD patterns were collected from 5° to 90° at a scanning speed of 10° min^−1^. Oriented samples of the clay fraction were scanned from 5° to 20° at the same speed. Ethylene glycol solvation and thermal treatment (500 °C for 1 h) were applied to the clay fraction and selected reference minerals to identify phyllosilicate phases. Reference minerals included enstatite, diopside (N’s Mineral Co. Ltd., Niigata, Japan), quartz (FUJIFILM Wako, Osaka, Japan), phlogopite (N’s Mineral Co. Ltd., Niigata, Japan), talc (Crown Talc PP JP Grade; Matsumura Sangyo Co. Ltd., Osaka, Japan), clinochlore, and a JCSS reference hydrobiotite (JCSS-5501; ((K_0.55_Na_0.08_Ca_0.06_)[Mg_2.34_Fe^3+^_0.39_Fe^2+^_0.08_Ti_0.07_][Si_3.02_Al_0.96_Fe^3+^_0.02_]O_10_(OH)_2_)^66^), which is an interstratified phlogopite/vermiculite mineral.

The 3-mm-thick rock section used for microbial characterizations was also examined by environmental scanning electron microscopy coupled with energy-dispersive X-ray spectroscopy (ESEM-EDS) without applying a conductive coating. ESEM-EDS analysis was done with a JSM-6510LA ESEM (JEOL, Tokyo, Japan) equipped with an EDS system, operated at 15-kV accelerating voltage and 60–80-Pa chamber pressure. The raw X-ray intensities were collected with an acquisition time of 60 seconds and processed using the ZAF correction method via the JEOL Analysis Station software. For semi-quantitative analysis, standardless-based procedures were applied.

μ-XRF mapping was performed using an SFXM at the beamline BL17SU at the SPring-8. At the BL17SU, the incident soft X-ray beam was focused to ca. 1.5 μm (vertical) × 3.5 μm (horizontal) using a Wolter mirror as a focusing component, and the fluorescent X-rays emitted from the sample were detected with a silicon drift detector in the PFY mode^67–69^. An excitation energy of 2000 eV was applied for the elemental mapping of Al (Fig. 2b). After μ-XRF mappings, Al *K*-edge, Fe *L_3_*-edge, and S *K*-edge XANES spectra were collected from the rock section and reference materials, including phlogopite, hydrobiotite, talc, diopside, hematite (FUJIFILM Wako, Osaka, Japan), elemental sulfur (FUJIFILM Wako, Osaka, Japan), sodium sulfite (Na_2_SO_3_; FUJIFILM Wako, Osaka, Japan), and sodium sulfate (Na_2_SO_4_; FUJIFILM Wako, Osaka, Japan). Excitation energy ranges for the Al *K*-edge, Fe *L_3_*-edge, and S *K*-edge were ca. 1550–1600 eV, 700–715 eV, and 2460–2490 eV with energy steps of 0.47 eV,0.15 eV, and 0.22 eV, respectively. The S *K*-edge XANES spectra of troilite, marcasite, and pyrite were obtained from the ID21 Sulfur XANES spectra database at the European Synchrotron Radiation Facility (ESRF)^70^. The XANES spectra were calibrated and normalized using the Athena software^71^.

To determine the iron valence state without being exposed to the atmosphere, we prepared a new 3-mm-thick rock section using a precision diamond wire saw (DWS 3400; Meiwa Fosis Corp., Tokyo, Japan) inside an Ar-purged glove box. The rock section was loaded into a He-purged sample holder at the beamline BL36XU of the SPring-8 for hard X-ray SFXM analysis. Oxygen levels in the glove box and the sample holder were less than 0.5 ppm. The incident hard X-ray beam was focused to ca. 1.6 μm (vertical) × 0.3 μm (horizontal) with a KB mirror as a focusing component, and the fluorescent X-rays emitted from the sample were detected with a silicon drift detector (Vortex-ME4; Hitachi High-Tech Corp., Tokyo, Japan) in the PFY mode^72^. An excitation energy of 8000 eV was applied for the elemental mapping of K. After μ-XRF mappings, Fe *K*-edge XANES spectra were collected from the rock section and nontronite (NAu-2; ((M^+^_0.97_)[Si_7.57_Al_0.01_Fe_0.42_][Al_0.52_Fe_3.32_Mg_0.7_]O_20_(OH)_4_)^28,73^ (Fe(III) standard). An excitation energy range for the Fe *K*-edge was 7105–7120 eV with an energy step of 0.35eV. The Fe *K*-edge XANES spectrum of FeCl_2_ (Fe(II) standard) was obtained from the XAFS Standard Sample Database at BL14B2 in SPring-8^29^. The Fe *K*-edge XANES spectra were also calibrated and normalized using the Athena software.

To assess chemical homogeneity, reference minerals without external certification (enstatite, diopside, talc, phlogopite, and clinochlore) were analyzed using an ESEM-EDS system under high-vacuum conditions (Extended Data Fig. 6). SEM specimens were prepared by embedding the samples in LR White resin, followed by polishing and carbon coating.

**Extended Data Fig. 6.**
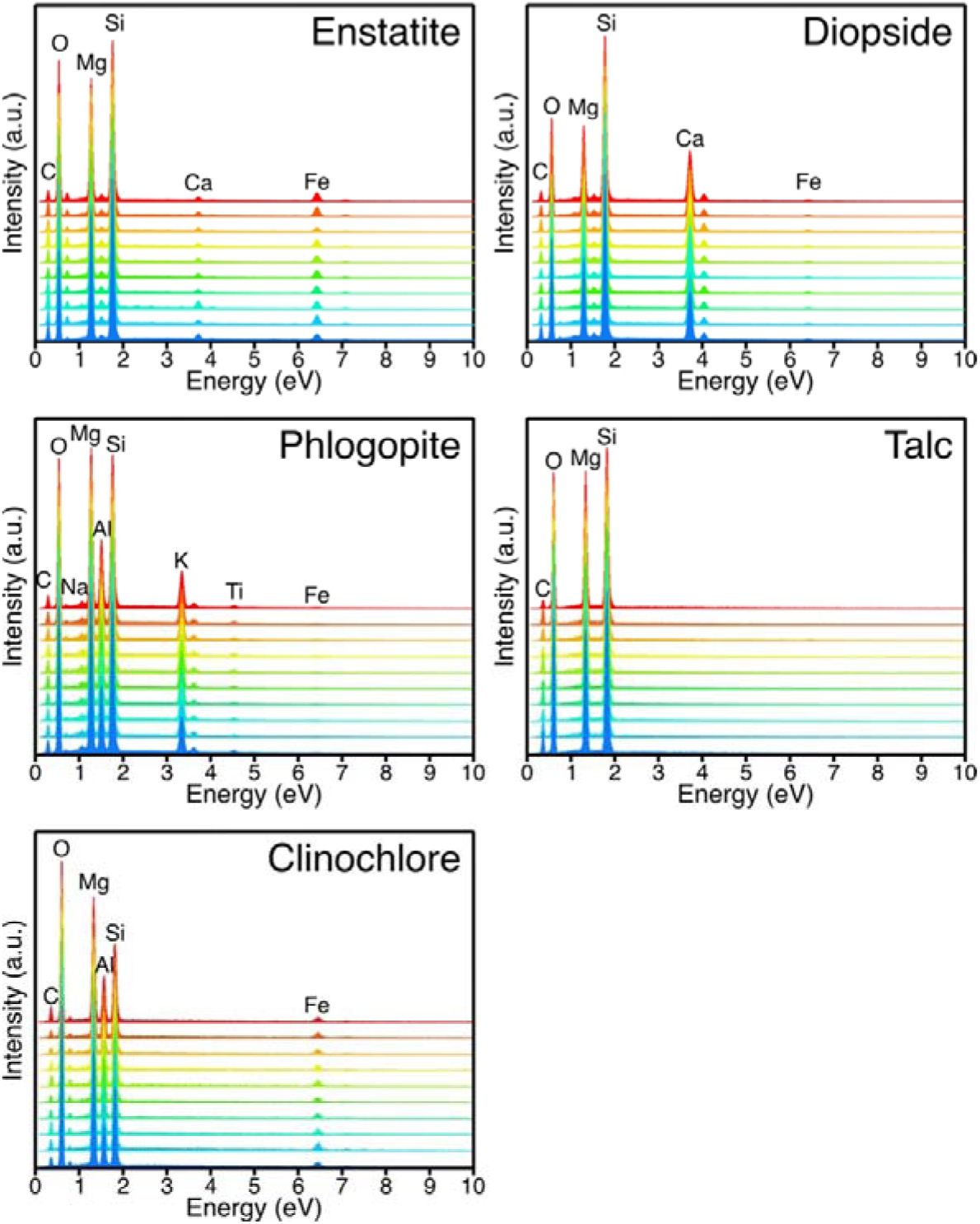
Chemical homogeneity of reference minerals. Stacked energy-dispersive X-ray spectroscopy (EDS) spectra from 10 grains each of enstatite, diopside, phlogopite, talc, and clinochlore demonstrate compositional homogeneity. Peak intensities for major elements are consistent with the expected stoichiometry of each mineral.

## Data availability

All data needed to evaluate the conclusions in the paper are present in the paper and/or the Supplementary Materials.

## Acknowledgments

We are grateful to Master Drilling for their support during the drilling operations. The authors would like to acknowledge Mpho Molautsi, Katja Heeschen, and Kwena Mathopa for their on-site assistance. We thank Eoghan Dillon (Photothermal Spectroscopy Corp.) for insightful discussions and technical advice regarding O-PTIR measurements. We also thank the beamline scientists at NanoTerasu (BL13U) and SPring-8 (BL17SU and BL36XU) for their technical support.

μ-XRF and XANES measurements were performed at BL13U of NanoTerasu with the approval of the Japan Synchrotron Research Institute (JASRI) (Proposal No. 2025A9013), and at BL17SU and BL36XU of SPring-8 with the approval of RIKEN (Proposal No. 20250069) and JASRI (Proposal No. 2025A1617). ESEM-EDS analysis was supported by the “Advanced Research Infrastructure for Materials and Nanotechnology in Japan (ARIM)” of the Ministry of Education, Culture, Sports, Science and Technology (MEXT) (Proposal Nos. JPMXP1224UT0317 and JPMXP1225UT0023).

The Bushveld Drilling Project (BVDP) was supported by the International Continental Scientific Drilling Program, the National Research Foundation of South Africa, the Deutsche Forschungsgemeinschaft (DFG) (Grant No. 684792 to J.K.), and the South African Council for Geoscience. This work was also supported by the JSPS/NRF Bilateral Joint Research Project (Grant No. JPJSBP120246501 to Y.S. and J.C.), the Astrobiology Center Program of National Institutes of Natural Sciences (NINS) (Grant No. AB0502 to Y.S.), JSPS KAKENHI (Grant Nos. JP25K22489 to Y.S. and JP25KJ1046 to T.K.), and JST SPRING (Grant No. JPMJSP2108 to T.K.). The author would like to thank Enago (www.enago.jp) for the English language review.

## Author Contributions

F.R., R.K., L.A., S.W., and R.T. contributed to the successful application and implementation of the Bushveld ICDP project. J.C., J.K., K.M., S.H., A.A., S.W., T.K., and Y.S. prepared and performed on-site sample collection. J.K. planned and set up the on-site contamination control, and C.N., T.M., F.R., and M.M. performed the geological curation of the examined rock sample. T.K., M.K., and Y.S. conducted the experimental work, including data collection and analysis. H.K. assisted with O-PTIR analysis. H.S., T.K., and T.W. assisted with synchrotron-based experiments at NanoTerasu. H.S., T.I., T.K., T.U, and M.O. assisted with synchrotron-based experiments at SPring-8. F.R., R.K., L.A., R.T., T.K., and Y.S. drafted the manuscript. All authors discussed the results and approved the final manuscript.

## Competing Interest Declaration

All authors declare that they have no competing interests.

## Notes

### Competing Interest Statement

The authors have declared no competing interest.

## References

1 Warr, O. et al. 86Kr excess and other noble gases identify a billion-year-old radiogenically-enriched groundwater system. Nat. Commun. 13, 3768 (2022).

2 Holland, G., Lollar, B. S., Li, L., Lacrampe-Couloume, G., Slater, G. F. & Ballentine, C. J. Deep fracture fluids isolated in the crust since the Precambrian era. Nature 497, 357–360 (2013).

3 Onstott, T. C. et al. Paleo–rock-hosted life on Earth and the search on Mars: a review and strategy for exploration. Astrobiology 19, 1230–1262 (2019).

4 O’Driscoll, B. & VanTongeren, J. A. Layered intrusions: from petrological paradigms to precious metal repositories. Elements 13, 383–389 (2017).

5 Cawthorn, R. (2015). The Bushveld Complex, South Africa. In: Charlier, B., Namur, O., Latypov, R., Tegner, C. (eds) Layered Intrusions. Springer Geology. Springer, Dordrecht.

6 Suzuki, Y., Webb, S. J., Kouduka, M. et al. Subsurface microbial colonization at mineral-filled veins in 2-billion-year-old mafic rock from the Bushveld Igneous Complex, South Africa. Microb. Ecol. 87, 116 (2024).

7 Suzuki, Y. et al. Deep microbial proliferation at the basalt interface in 33.5–104 million-year-old oceanic crust. Commun. Biol. 3, 131 (2020).

8 Zema, M., Ventruti, G., Iacalamita, M. & Scordari, F. Kinetics of Fe-oxidation/deprotonation process in Fe-rich phlogopite under isothermal conditions. Am. Mineral. 95, 1458–1466 (2010).

9 Ma, T., Sun, H., Peng, T. & Zhang, Q. Transformation process from phlogopite to vermiculite under hydrothermal conditions. Appl. Clay Sci. 208, 106094 (2021).

10 Francis, D. Columbia Hills—an exhumed layered igneous intrusion on Mars? Earth Planet. Sci. Lett. 310, 59–64 (2011).

11 Bedle, H., Cooper, C. M. & Frost, C. D. Nature versus nurture: preservation and destruction of Archean cratons. Tectonics 40, e2021TC006714 (2021).

12 Nisson, D. M. et al. Radiolytically reworked Archean organic matter in a habitable deep ancient high-temperature brine. Nat. Commun. 14, 6163 (2023).

13 Sherwood Lollar, B., et al. A window into the abiotic carbon cycle: acetate and formate in fracture waters in 2.7 billion year-old host rocks of the Canadian Shield. Geochim. Cosmochim. Acta 294, 295–314 (2021).

14 Nisson, D. M. et al. Hydrogeochemical and isotopic signatures elucidate deep subsurface hypersaline brine formation through radiolysis-driven water–rock interaction. Geochim. Cosmochim. Acta 340, 65–84 (2023).

15 Lollar, G. S. et al. ‘Follow the water’: Hydrogeochemical constraints on microbial investigations 2.4 km below surface at the Kidd Creek Deep Fluid and Deep Life Observatory. Geomicrobiol. J. 36, 859–872 (2019).

16 Hawkesworth, C., Cawood, P. A., Dhuime, B. & Kemp, T. Tectonic processes and the evolution of the continental crust. J. Geol. Soc. 181, jgs2024-027 (2024).

17 Becker, M., Harris, P. J., Wiese, J. G. & Bradshaw, D. J. Mineralogical characterisation of naturally floatable gangue in Merensky Reef ore flotation. Int. J. Miner. Process. 93, 246–255 (2009).

18 Buick, I. S., Maas, R. & Gibson, R. Precise U–Pb titanite age constraints on the emplacement of the Bushveld Complex, South Africa. J. Geol. Soc. 158, 3–6 (2001).

19 Allwright, A. J., de Lange, S., Lubbe, R., Mbonambi, L., Vivier, K. & Witthuser, K. T. Research-based exploration of deep groundwater within the eastern limb of the Bushveld Igneous Complex for hydrogeological characterisation and potential future water resource identification. Final report. Water Research Commission, Pretoria (2025). ISBN 978-0-6392-0706-3.

20 Friese, A. et al. A simple and inexpensive technique for assessing contamination during drilling operations. Limnol. Oceanogr. Methods, 15, 200–211 (2017).

21 Leinweber, P., Kruse, J., Walley, F. L., Gillespie, A., Eckhardt, K. U., Blyth, R. I. & Regier, T. Nitrogen K-edge XANES—an overview of reference compounds used to identify ‘unknown’ organic nitrogen in environmental samples. J. Synchrotron Radiat. 14, 500–511 (2007).

22 Gillespie, A. W., Walley, F. L., Farrell, R. E., Regier, T. Z. & Blyth, R. I. Calibration method at the N K-edge using interstitial nitrogen gas in solid-state nitrogen-containing inorganic compounds. J. Synchrotron Radiat. 15, 532–534 (2008).

23 Moore, D. M. & Reynolds, R. C. Jr. X-ray diffraction and the identification and analysis of clay minerals, 2nd edn (Oxford Univ. Press, 1997).

24 Li, D., Bancroft, G. M., Fleet, M. E., Feng, X.-H. & Pan, Y. Al *K*-edge XANES spectra of aluminosilicate minerals. Am. Mineral. 80, 432–440 (1995).

25 van Aken, P. A. & Liebscher, B. Quantification of ferrous/ferric ratios in minerals: new evaluation schemes of Fe L₂,₃ electron energy-loss near-edge spectra. Phys. Chem. Miner. 29, 188–200 (2002).

26 Van Groeningen, N., ThomasArrigo, L. K., Byrne, J. M., Kappler, A., Christl, I. & Kretzschmar, R. Interactions of ferrous iron with clay mineral surfaces during sorption and subsequent oxidation. Environ. Sci. Process. Impacts 22, 1355–1367 (2020).

27 Berry, A. J., O’Neill, H. S. C., Jayasuriya, K. D., Campbell, S. J. & Foran, G. J. XANES calibrations for the oxidation state of iron in a silicate glass. Am. Mineral. 88, 967–977 (2003).

28 Qian, Y., Scheinost, A. C., Grangeon, S., Greneche, J.-M., Hoving, A., Bourhis, E., Maubec, N., Churakov, S. V. & Fernandes, M. M. Oxidation state and structure of Fe in nontronite: from oxidizing to reducing conditions. ACS Earth Space Chem. 7, 1868–1881 (2023).

29 Ofuchi, H., Matsumoto, T. & Honma, T. Construction of XAFS standard sample database at BL14B2 in SPring-8. Radiat. Phys. Chem. 218, 111581 (2024).

30 Hogg, C. S. & Meads, R. E. A Mössbauer study of thermal decomposition of biotites. Mineral. Mag. 40, 79–88 (1975).

31 Cawthorn, R. G. & Walraven, F. Emplacement and crystallization time for the Bushveld Complex. J. Petrol. 39, 1669–1687 (1998).

32 Li, C., Ripley, E. M., Sarkar, A., Shin, D. & Maier, W. D. Origin of phlogopite-orthopyroxene inclusions in chromites from the Merensky Reef of the Bushveld Complex, South Africa. Contrib. Mineral. Petrol. 150, 119–130 (2005).

33 Wagner, W. & Pruß, A. The IAPWS formulation 1995 for the thermodynamic properties of ordinary water substance for general and scientific use. J. Phys. Chem. Ref. Data 31, 387–535 (2002).

34 Scoates, J. S., Wall, C. J., Friedman, R. M., Weis, D., Mathez, E. A. & VanTongeren, J. A. Dating the Bushveld Complex: timing of crystallization, duration of magmatism, and cooling of the world’s largest layered intrusion and related rocks. J. Petrol. 62, 1–39 (2021).

35 de la Calle, C. & Suquet, H. Vermiculite. In Hydrous Phyllosilicates (Exclusive of Micas) (ed. Bailey, S. W.) 455–496 (De Gruyter, 1988).

36 Janzen, M. P., Nicholson, R. V. & Scharer, J. M. Pyrrhotite reaction kinetics: reaction rates for oxidation by oxygen, ferric iron, and for nonoxidative dissolution. Geochim. Cosmochim. Acta 64, 1511–1522 (2000).

37 Beaver, R. C. & Neufeld, J. D. Microbial ecology of the deep terrestrial subsurface. ISME J. 18, wrae091 (2024).

38 Dong, H., Zeng, Q., Sheng, Y. et al. Coupled iron cycling and organic matter transformation across redox interfaces. Nat. Rev. Earth Environ. 4, 659–673 (2023).

39 Price, P. B. & Sowers, T. Temperature dependence of metabolic rates for microbial growth, maintenance, and survival. Proc. Natl Acad. Sci. USA 101, 4631–4636 (2004).

40 Hoehler, T. & Jørgensen, B. Microbial life under extreme energy limitation. Nat. Rev. Microbiol. 11, 83–94 (2013).

41 Templeton, A. S. & Caro, T. A. The rock-hosted biosphere. Annu. Rev. Earth Planet. Sci. 51, 493–519 (2023).

42 do Nascimento Vieira, A., Kleinermanns, K., Martin, W. F. & Preiner, M. The ambivalent role of water at the origins of life. FEBS Lett. 594, 2717–2733 (2020).

43 Ménez, B., Pasini, V. & Brunelli, D. Life in the hydrated suboceanic mantle. Nat. Geosci. 5, 133–137 (2012).

44 Klein, F., Humphris, S. E., Guo, W., Schubotz, F., Schwarzenbach, E. M. & Orsi, W. D. Fluid mixing and the deep biosphere of a fossil Lost City-type hydrothermal system at the Iberia Margin. Proc. Natl Acad. Sci. USA 112, 12036–12041 (2015).

45 Lima-Zaloumis, J., Neubeck, A., Ivarsson, M., Bose, M., Greenberger, R., Templeton, A. S. et al. Microbial biosignature preservation in carbonated serpentine from the Samail Ophiolite, Oman. Commun. Earth Environ. 3, 231 (2022).

46 Ivarsson, M., Bach, W., Broman, C., Neubeck, A. & Bengtson, S. Fossilized life in subseafloor ultramafic rocks. Geomicrobiol. J. 35, 460–467 (2018).

47 Suzuki, Y., Koduka, M., Brenker, F. E., Brooks, T., Glamoclija, M., Graham, H. V. et al. Submicron-scale detection of microbes and smectite from the interior of a Mars-analogue basalt sample by optical-photothermal infrared spectroscopy. Int. J. Astrobiol. 24, e1 (2025).

48 Li, X., Zhang, D., Bai, Y., Wang, W., Liang, J. & Cheng, J. X. Fingerprinting a living cell by Raman integrated mid-infrared photothermal microscopy. Anal. Chem. 91, 10750–10756 (2019).

49 Ménez, B. et al. Abiotic synthesis of amino acids in the recesses of the oceanic lithosphere. Nature 564, 59–63 (2018).

50 Hickman-Lewis, K., Cuadros, J., Yi, K. et al. Aluminous phyllosilicates promote exceptional nanoscale preservation of biogeochemical heterogeneities in Archaean siliciclastic microbial mats. Nat. Commun. 16, 2726 (2025).

51 Tokumaru, A. et al. Co-enrichment of Ce and organics in microbe-like structures at the deep-sea ferromanganese crust surface. Phil. Trans. R. Soc. A 383, 20240432 (2026).

52 Chan, C. S. et al. Lithotrophic iron-oxidizing bacteria produce organic stalks to control mineral growth: implications for biosignature formation. ISME J. 5, 717–727 (2011).

53 Suga, H. et al. Spatially resolved distribution of Fe species around microbes at the submicron scale in natural bacteriogenic iron oxides. Microbes Environ. 32, 283–287 (2017).

54 Attwood, D. Soft X-rays and extreme ultraviolet radiation: principles and applications. Cambridge Univ. Press (2000).

55 Ming, D. W., Mittlefehldt, D. W., Morris, R. V., Golden, D. C., Gellert, R., Yen, A. et al. Geochemical and mineralogical indicators for aqueous processes in the Columbia Hills of Gusev crater, Mars. J. Geophys. Res. Planets 111, E02S12 (2006).

56 Liu, Y., Tice, M. M., Schmidt, M. E., Treiman, A. H., Kizovski, T. V., Hurowitz, J. A., Zorzano, M. P. et al. An olivine cumulate outcrop on the floor of Jezero crater, Mars. Science 377, 1513–1519 (2022).

57 Farley, K. A., Stack, K. M., Shuster, D. L., Horgan, B. H. N., Hurowitz, J. A., Tarnas, J. D. et al. Aqueously altered igneous rocks sampled on the floor of Jezero crater, Mars. Science 377, eabo2196 (2022).

58 Hurowitz, J. A., Tice, M. M., Allwood, A. C., Cable, M. L., Hand, K. P., Murphy, A. E., et al. Redox-driven mineral and organic associations in Jezero Crater, Mars. Nature 645, 332–340 (2025).

59 Hollis, J. R., Abbey, W., Beegle, L. W., Bhartia, R., Ehlmann, B. L., Miura, J., Monacelli, B., Moore, K., Nordman, A. & Scheller, E. A deep-ultraviolet Raman and fluorescence spectral library of 62 minerals for the SHERLOC instrument onboard Mars 2020. Planet. Space Sci. 209, 105356 (2021).

60 Barnes, J., McIntyre, J. R., Nisbet, B. W. & Williams, C. R. Platinum-group element mineralisation in the Munni Munni Complex, Western Australia. Mineral. Petrol. 42, 141–164 (1990).

61 Barnes, S. J. & Hoatson, D. M. The Munni Munni complex, Western Australia: stratigraphy, structure and petrogenesis. J. Petrol. 35, 715–751 (1994).

62 Mukasa, S. B., Wilson, A. H. & Carlson, R. W. A multielement geochronologic study of the Great Dyke, Zimbabwe: significance of the robust and reset ages. Earth Planet. Sci. Lett. 164, 353–369 (1998).

63 Wilson, A. H. & Prendergast, M. D. Platinum-group element mineralisation in the Great Dyke, Zimbabwe, and its relationship to magma evolution and magma chamber structure. S. Afr. J. Geol. 104, 319–342 (2001).

64 Järvinen, V., Halkoaho, T., Konnunaho, J., Heinonen, J. S. & Rämö, O. T. Parental magma, magmatic stratigraphy and reef-type PGE enrichment of the 2.44 Ga mafic–ultramafic Näränkävaara layered intrusion, northern Finland. Mineralium Deposita 55, 1535–1560 (2020).

65 Becraft, E. D. et al. Evolutionary stasis of a deep subsurface microbial lineage. ISME J. 15, 2830–2842 (2021).

66 Kikuchi, R. & Kogure, T. Structural and compositional variances in ‘hydrobiotite’ sample from Palabora, South Africa. Clay Sci. 22, 39–52 (2018).

67 Oura, M., Suga, H., & Suzuki, Y. Introduction to scanning fluorescence X-ray microscopy and its applications for earth and planetary science. Jpn. Mag. Mineral. Petrol. Sci. 55, gkk.250920 (2026). (in Japanese).

68 SPring-8·SACLA Annual Report FY2023, 99–102.

69 SPring-8·SACLA Annual Report FY2024, 99–103.

70 ID21 Sulfur XANES spectra database, European Synchrotron Radiation Facility (ESRF). Created by E. Chalmin and ID21 users. Available at: https://www.esrf.fr/home/UsersAndScience/Experiments/ID21/php.html (Accessed: 18 February 2026).

71 Ravel, B. & Newville, M. ATHENA, ARTEMIS, HEPHAESTUS: data analysis for X-ray absorption spectroscopy using IFEFFIT. J. Synchrotron Radiat. 12, 537–541 (2005).

72 Uruga, T., Tada, M., Sekizawa, O., Takagi, Y., Yokoyama, T. & Iwasawa, Y. SPring-8 BL36XU: Synchrotron radiation X-ray-based multi-analytical beamline for polymer electrolyte fuel cells under operating conditions. Chem. Rec. 19, 1444–1456 (2019).

73 Keeling, J. L., Raven, M. D., & Gates, W. P. Geology and characterization of two hydrothermal nontronites from weathered metamorphic rocks at the Uley graphite mine, South Australia. Clays Clay Miner. 48, 537–548 (2000).

